# Efficient RNP-directed human gene targeting reveals SPDEF is required for IL-13-induced mucostasis

**DOI:** 10.1101/716670

**Authors:** Kyung Duk Koh, Sana Siddiqui, Dan Cheng, Luke R. Bonser, Dingyuan I. Sun, Lorna T. Zlock, Walter E. Finkbeiner, Prescott G. Woodruff, David J. Erle

## Abstract

Primary human bronchial epithelial cell (HBEC) cultures are a useful model for studies of lung health and major airway diseases. However, mechanistic studies have been limited by our ability to selectively disrupt specific genes in these cells. Here we optimize methods for gene targeting in HBECs by direct delivery of single guide RNA (sgRNA) and recombinant Cas9 (rCas9) complexes by electroporation, without a requirement for plasmids, viruses, or antibiotic selection. Variations in the method of delivery, sgRNA and rCas9 concentrations, and sgRNA sequences all had effects on targeting efficiency, allowing for predictable control of the extent of gene targeting and for near complete disruption of gene expression. To demonstrate the value of this system, we targeted *SPDEF*, which encodes a transcription factor previously shown to be essential for the differentiation of MUC5AC-producing goblet cells in mouse models of asthma. Targeting *SPDEF* led to proportional decreases in *MUC5AC* expression in HBECs stimulated with IL-13, a central mediator of allergic asthma. Near-complete targeting of *SPDEF* abolished IL-13-induced *MUC5AC* expression and goblet cell differentiation. In addition, targeting of *SPDEF* prevented IL-13-induced impairment of mucociliary clearance, which is likely to be an important contributor to airway obstruction, morbidity, and mortality in asthma. We conclude that direct delivery of sgRNA and rCas9 complexes allows for predictable and efficient gene targeting and enables mechanistic studies of disease-relevant pathways in primary HBECs.

## Introduction

The bronchial epithelium is critical for normal lung function, and altered epithelial function is central to the pathogenesis of major airway diseases including asthma, chronic obstructive pulmonary disease, and cystic fibrosis (1, 2). The mechanisms underlying normal bronchial epithelial cell function and pathophysiological alterations leading to disease have been studied in various model systems. Mouse and other animal model systems have made indispensable contributions to our understanding of airway epithelial cells, but intra-species differences (3) require that we complement these studies with others in human systems. *In vivo* investigations in humans are important but are largely confined to observational studies until pre-clinical work justifies clinical trials of therapeutic agents. Studies in immortalized cell lines are also valuable, but these systems are quite limited in their ability to model differentiation of ciliated cells, mucus-producing goblet cells, and other specialized airway epithelial subpopulations with critical roles in human health and disease (4). Primary air-liquid interface (ALI) cultures of human bronchial epithelial cells (HBECs) have proven to be valuable models due to their ability to recapitulate many *in vivo* aspects of airway epithelial cell differentiation and function of the human airway epithelium (5, 6). Whereas gene targeting has enabled elegant mechanistic studies in transgenic mice, analyses of the contributions of specific genes in highly differentiated HBEC models have been limited due to challenges in efficiently targeting genes in these cells.

The advent of CRISPR/Cas9 gene editing technology has transformed our ability to investigate gene function in many model systems and has great potential for therapeutic applications. CRISPR/Cas9 gene editing makes use of a programmable guide RNA (gRNA) as well as a Cas9 nuclease to form a ribonucleoprotein (RNP) complex, which in turn binds and cleaves the target site in genomic DNA (7, 8). gRNA and Cas9 have been delivered to human cells using viral and non-viral delivery systems (9). Viral delivery of gRNA and Cas9 transgenes together with an antibiotic resistance gene has been used for gene targeting in many cell types, including HBECs (10). Viral delivery allows for selection of transduced cells, which improves efficiency but requires the use of more starting cells or more rounds of cell division. Moreover, continued expression of Cas9 may increase the likelihood of undesired effects, including chromosome rearrangements (11). An alternative to viral delivery is direct delivery of ribonucleoprotein (RNP) complexes of guide RNA (gRNA) and recombinant Cas9 (rCas9) via electroporation. This approach has been shown to target genes at high efficiency in many cell types, including primary cells (12–14).

Here, we describe methods for direct delivery of RNP complexes to robustly target the genome of primary HBECs. As an illustration of the value of these methods, we targeted the transcription factor *SPDEF.* Previous studies with *Spdef*^*-/-*^ transgenic mice demonstrated that this transcription factor is required for airway epithelial cell mucus metaplasia in mouse models of asthma (15). *Spdef* is induced by IL-13, a key mediator in type 2 asthma, and regulates the expression of a substantial set of transcripts found in mucus-producing goblet cells. By targeting *SPDEF* in HBECs, we were able to examine its contributions to IL-13-induced changes in gene expression and epithelial function in primary human cells.

## Materials and Methods

### CRISPR-based gene targeting

gRNA sequences were selected from the CRISPR Targets 10K track on the UCSC Genome Browser (https://genome.ucsc.edu, hg38 human genome build) and are shown in Table E1. RNAs were resuspended in 150 mM KCl and 10 mM Tris-HCl, pH 7.4. For experiments involving crRNA and tracrRNA (Thermo Fisher Scientific, Waltham, MA), crRNA (160 *µ*M, 1 *µ*L) and tracrRNA (160 *µ*M, 1 *µ*L) were mixed, hybridized at 37 °C for 30 min, combined with rCas9 (40 *µ*M, 2 *µ*L; MacroLab, Berkeley, CA) and electroporation enhancer DNA oligonucleotide (100 *µ*M, 1 *µ*L), and incubated at 37 °C for 15 min prior to electroporation. Unless otherwise indicated, for experiments involving sgRNAs (Synthego, Redwood City, CA), sgRNA (53.3 *µ*M, 3 *µ*L) was combined with rCas9 (40 *µ*M, 2 *µ*L), and incubated at room temperature for 10 min prior to electroporation.

After reaching 80% confluence, HBECs or BEAS-2B cells were harvested for electroporation. Unless otherwise indicated, 150,000 cells were resuspended in 20 *µ*L of P3 Primary Cell Nucleofector Solution with Supplement 1 and mixed with the gRNA/rCas9 solutions before electroporation (4D-Nucleofector System, Lonza, Walkersville, MD; program DC-100). For experiments involving the Neon Transfection System (Thermo Fisher Scientific), ∼150,000 cells were resuspended in Resuspension Buffer R, mixed with the gRNA/rCas9 solutions, and electroporated at 1400 V, 20 ms, and 2 pulses. For experiments involving two sequential electroporations, HBECs were seeded on human placental collagen (HPC)-coated 6-well dishes and propagated in BEGM supplemented with the rho-associated protein kinase inhibitor Y-27632 (10 *µ*M; Enzo Life Sciences, Farmingdale, NY), which enhances cell proliferation (16). After 3 d, cells were harvested for the second electroporation. After completing the electroporation(s), HBECs were seeded on HPC-coated 6.5 mm Transwell inserts (Corning, Corning, NY) and cultured at ALI as described previously (17). HBECs were maintained in ALI culture for 23 d before harvesting. Where indicated, IL-13 (10 ng/mL; Peprotech, Rocky Hill, NJ) was added to the culture medium for the final 7 d of culture. In experiments with BEAS-2B cells, cells were harvested 3 d after electroporation.

### Measuring targeting efficiency

Genomic DNA was extracted using the Quick-DNA Miniprep Plus Kit (Zymo Research, Irvine, CA) or the RNA/DNA/Protein Purification Plus Kit (Norgen Biotek, Thorold, ON, Canada) according to manufacturers’ protocols. ∼1 kb regions containing the gRNA target were amplified using appropriate PCR primers (Table E2) and Q5 DNA Polymerase (New England Biolabs, Ipswich, MA) and purified using a PCR Purification Kit (Qiagen, Germantown, MD). In cases where one gRNA was used per sample, PCR products were analyzed by Sanger sequencing (MCLAB, South San Francisco, CA; sequencing primers are shown in Table E2), and sequencing read files from cells treated with gene targeting gRNAs (experimental) and with non-targeting gRNAs (control) were analyzed using the Inference of CRISPR Edits (ICE) tool (18) to determine targeting efficiency, defined as percentage of insertions or deletions (indels). In experiments involving deletion of a larger genomic DNA region using a pair of sgRNAs, targeting efficiency was measured via 2% agarose gel electrophoresis followed by quantitation of PCR product bands intensities using Fiji (19). Band intensities were adjusted according to the estimated masses of each PCR product.

Additional information regarding materials and methods can be found in the Online Data Supplement.

## Results

### Optimizing genome editing in human primary bronchial epithelial cells

Our general approach to gene editing in HBECs via electroporation of gRNA/rCas9 RNP complex is shown in Figure 1A. After 6 d in submerged culture, cells were placed in suspension and electroporated using recombinant Cas9 enzyme (rCas9) complexed with a CRISPR RNA (crRNA) designed to target *SPDEF* and a trans-activating CRISPR RNA (tracrRNA). Cells were then placed in culture on Transwell inserts. After reaching confluence (day 0), the apical medium was removed, and cells were maintained in ALI culture. Cells were harvested at d 3 and gene editing efficiency was analyzed by ICE-coupled Sanger sequencing (18). We began by using rCas9, crRNA, and tracrRNA concentrations previously used with primary human T cells (14). In initial experiments, we used the 4D-Nucleofector system and found similar editing efficiencies (∼30%) using each of 18 nucleofection programs (Figure 1B). Editing efficiency was similar when cell numbers were doubled (from 75,000 to 150,000 cells per electroporation). We compared the 4D-Nucleofector and Neon nucleofection systems and did not observe consistent differences in efficiency (Figure 1C). However, addition of a second round of electroporation, performed 3 d after the first electroporation, substantially increased efficiency. The 4D-Nucleofector system, which allows for up to 96 simultaneous electroporations, was used for subsequent experiments.

**Figure 1.**
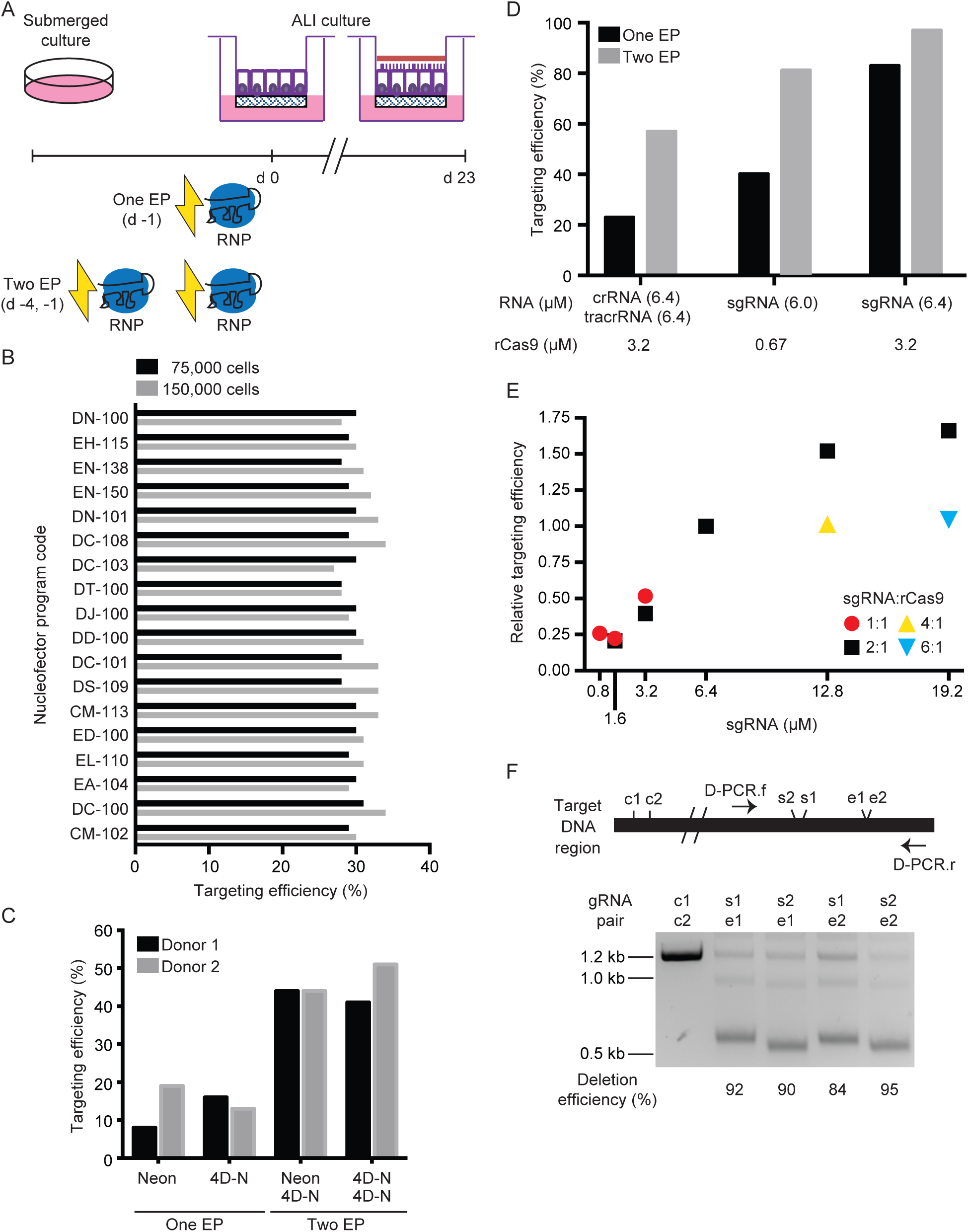
Optimization of gene targeting via RNP transfection of primary HBECs. (A) Overview of targeting protocol. Following a period of submerged culture on tissue culture dishes, cells were mixed with gRNA:rCas9 RNP complexes that were delivered by electroporation (EP) prior to initiation of ALI culture. In some experiments, cells underwent two EP prior to initiation of ALI. Cells were cultured at ALI for 23 d prior to harvesting. (B) Effects of 4D-Nucleofector program on targeting efficiency. RNP complexes comprised of SPDEF-3 crRNA (6.4 *µ*M), tracrRNA (6.4 *µ*M), and rCas9 (3.2 *µ*M) were delivered to either 75,000 (black) or 150,000 (grey) primary HBECs using 18 different programs. All subsequent experiments were performed with program DC-100 and 150,000 cells. (C) Effects of nucleofection device and number of EPs on targeting efficiency. RNP complexes were prepared as in (B) and delivered to HBECs using either a Neon Nucleofection System or a 4D-Nucleofector (4D-N). After one EP, cells were either directly seeded on Transwells or propagated in submerged culture for 3 d prior to undergoing a second EP. All subsequent experiments were performed using the 4D-N device. (D) Comparison of crRNA:tracrRNA and sgRNA-based gene targeting. The crRNA and the sgRNA were designed based on the same SPDEF-3 gRNA sequence and delivered by a single electroporation or by two sequential electroporations. (E) Effects of varying sgRNA and rCas9 concentrations. Results are aggregated from three separate single electroporation experiments. In one experiment, a KLF5 sgRNA was used to test sgRNA:rCas9 ratios of 4:1 and 6:1. In two other experiments, the SPDEF-3 sgRNA was used to test other ratios. In all experiments, one sample was electroporated with the reference concentrations (sgRNA 6.4 *µ*M, rCas9 3.2 *µ*M; 2:1 ratio). At the reference concentrations, absolute targeting efficiencies ranged from 55 to 70%. Relative targeting efficiencies were normalized to reference concentration targeting efficiencies from the same experiment to facilitate comparisons across experiments. The reference concentrations of sgRNA and rCas9 used in these experiments were used for all subsequent experiments. (F) Highly efficient deletion of a genomic DNA region. HBECs were transfected with 3.2 *µ*M rCas9 and a pair of sgRNAs (s1 or s2 and e1 or e2; 3.2 *µ*M of each of the pair of sgRNAs) by two electroporations, targeting two sites separated by ∼0.6 kb. Genomic DNA from targeted cells was amplified by PCR and analyzed by agarose gel electrophoresis. Control cells were transfected with c1 and c2 control sgRNAs, which targeted sequences located ∼15 kb outside of the amplified region.

In an attempt to improve editing efficiency, we focused on the type and amount of guide RNA. As an alternative to using the two component crRNA:tracrRNA system, we tested a single guide RNA (sgRNA) approach. Unlike the crRNA:tracrRNA system, which requires hybridization of two smaller RNA molecules that then complex with rCas9, each sgRNA molecule contains both the crRNA sequence and the Cas9-binding sequence. Compared with a crRNA:tracrRNA targeting the same sequence, delivery of sgRNA resulted in a higher editing efficiency (up to 97% with two electroporations, Figure 1D). When using sgRNA, increasing the amounts of RNA and rCas9 from levels recommended by the manufacturer (6.0 and 0.67 *µ*M, respectively) to the amount used in our standard crRNA:tracrRNA protocol (6.4 and 3.2 *µ*M) appeared to increase editing efficiency. To evaluate this in more detail, we tested a range of concentrations and ratios of sgRNA:rCas9 (Figure 1E). For a given concentration of rCas9, editing efficiency reached near maximal levels when sgRNA was present in a 2-fold molar excess. Efficiency increased when the amount of sgRNA was increased while maintaining a 2:1 ratio.

Strand breaks generated by targeting a single DNA sequence generally result in short indels, but for some applications, such as analysis of non-coding regions of the genome, it is useful to delete larger regions. To test whether our approach could be used to delete larger regions, we introduced pairs of sgRNAs targeting regions that were separated by ∼600 nt (Figure 1F). We found that this approach efficiently introduced large deletions in HBECs.

### Predictable control of gene targeting efficiency in HBECs

Editing efficiency also depends on the gRNA sequence. Despite the availability of numerous prediction algorithms, currently available computational tools cannot robustly predict gRNA editing efficiencies (20). Since the supply of HBECs can be a limiting factor for some experiments, we investigated whether HBEC editing efficiency could be estimated by testing gRNAs in BEAS-2B, an immortalized human airway epithelial cell line. Four different gRNAs targeting the same *SPDEF* exon were selected using the MIT Guide Design Tool, and all had high specificity scores. We delivered each to BEAS-2B cells (single electroporation) and primary HBE cells (single or double electroporation) (Figure 2A). The relative editing efficiency for the four gRNAs was the same in all cases, with gRNA 2 having the lowest efficiency and gRNA 4 the highest. For each sgRNA, editing efficiency in BEAS-2B cells (one electroporation) was similar to editing efficiency found after two electroporations of HBECs.

**Figure 2.**
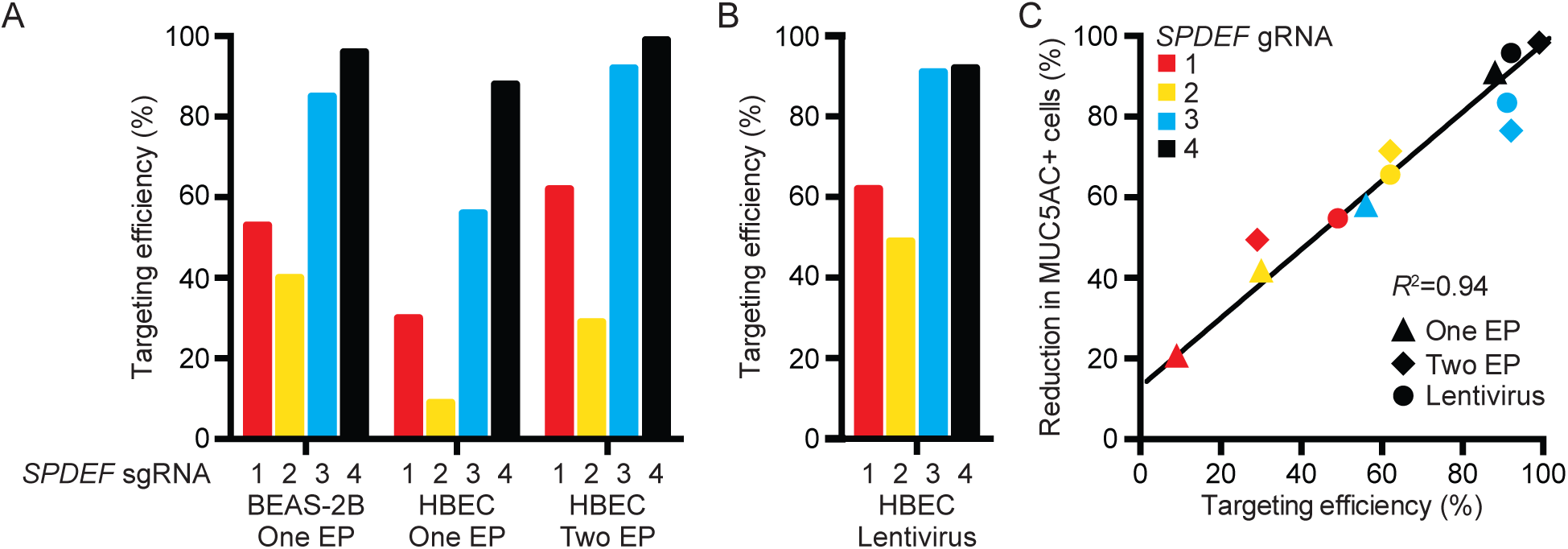
Predictable control of gene targeting efficiencies in primary HBECs. (A) Efficiencies of four sgRNAs targeting sequences in the *SPDEF* gene delivered to BEAS-2B cells or HBECs by electroporation. (B) Targeting efficiencies of the same gRNAs delivered to HBECs by lentiviral transduction. (C) After electroporation or viral transduction with same *SPDEF* targeting sgRNAs, HBEC ALI cultures were differentiated and stimulated with IL-13, and MUC5AC-producing cells were quantified by flow cytometry. Reduction in MUC5AC-producing cells was computed by comparison with cells transfected with non-targeting control gRNAs (NT) in the same experiment. MUC5AC measurements are paired with targeting efficiency measurements from duplicate wells of cells from the same transfection or transduction. Data are pooled from two experiments with different HBEC donors. *R*, Pearson correlation coefficient.

Although virus-free gene targeting offers many advantages, virus-based methods are also useful for some applications. Direct determination of virus-based gene targeting efficiency in HBECs requires cloning, virus production, and testing in cell culture. To determine whether the efficiency of virus-based targeting in HBECs could be predicted from results with virus-free targeting with RNPs, we produced lentiviruses that drove expression of each of the four *SPDEF* gRNAs. Lentiviral delivery of each of these four gRNAs followed by antibiotic selection for transduced cells (Figure 2B) produced editing efficiencies that were remarkably similar to efficiencies obtained with the same sequences delivered using the virus-free system without antibiotic selection (Figure 2A). This suggests that sequence-dependent effects on editing efficiency are largely independent of method of delivery, and that the virus-free methods can be useful for rapid screening of gRNA efficiency prior to subsequent experiments using either virus-free or virus-based methods.

In most cases, the goal of gene targeting is to eliminate functional protein production by the targeted gene. We attempted to use commercially available antibodies against SPDEF but were unable to detect this protein in untargeted cells. As an alternative, we measured the effect of *SPDEF* targeting on SPDEF function. Previous studies in mice demonstrated that *Spdef* is required for IL-13-induced production of the airway mucin MUC5AC (15). We used both virus-free and lentiviral-based methods to target *SPDEF* in HBECs, allowed cells to differentiate at ALI, stimulated cells with IL-13 for 7 d, and then measured MUC5AC-producing cells by flow cytometry. We found excellent correlation between DNA editing efficiency and reduction in MUC5AC-producing cells (Figure 2C). This correlation was seen across four distinct gRNA sequences, providing strong evidence that the effect results from elimination of functional SPDEF protein and not from off-target effects. The most efficient gRNA sequence, SPDEF-4, resulted in >90% *SPDEF* targeting and >95% reduction in MUC5AC-producing cells with both virus-free and lentiviral delivery.

### Highly efficient gene targeting dissects the role of *SPDEF* in IL-13-induced effects on the HBEC transcriptome

Having established the capacity for efficient targeting of *SPDEF*, we used the HBEC ALI culture system to study downstream effects of sgRNA/rCas9-mediated non-viral deletion of this critical transcription factor. Analyses of effects of *SPDEF* targeting on mRNA transcripts showed that SPDEF is required for IL-13-induced increases of *MUC5AC* (Figure 3A). SPDEF was also required for IL-13-induced increases in the goblet cell transcription factor *FOXA3* mRNA (Figure 3B), consistent with prior work indicating that *Foxa3* is downstream of *Spdef* in mouse airway epithelial cells (15). In addition, SPDEF was required for induction of *CLCA1* (Figure 3C), which has been shown to trigger a MAPK13-dependent pathway responsible for mucus production (21). In addition to increases in these goblet cell transcripts, it is known that IL-13 also suppresses expression of the Club cell secretory protein gene, *SCGB1A1* (also known as *CCSP*), indicating that IL-13 causes a shift of secretory cells from a predominantly Club cell phenotype to a predominantly goblet cell phenotype. We found that the IL-13-induced decrease in *SCGB1A1*, like the IL-13-induced increase in goblet cell genes, was SPDEF*-*dependent (Figure 3D). In mouse airway epithelial cells, IL-13 stimulation increases expression of both major airway mucins, *Muc5ac* and *Muc5b* (22). In humans, however, IL-13 strongly induces *MUC5AC* but modestly suppresses *MUC5B* (23). The availability of a system for efficient targeting in HBECs allowed us to investigate the role of SPDEF in *MUC5B* regulation in human cells. Although IL-13-stimulated HBECs that were SPDEF-deficient had levels of *MUC5AC, FOXA3, CLCA1*, and *SCGB1A1* that were similar to those found in unstimulated cells, *MUC5B* expression was not normalized (Figure 3A-E). Instead, *SPDEF* targeting led to a further decrease in expression of *MUC5B* below the level seen with IL-13 treatment alone. This indicates that therapeutic approaches that target *SPDEF* in asthma are likely to inhibit expression of both *MUC5AC* and *MUC5B.*

**Figure 3.**
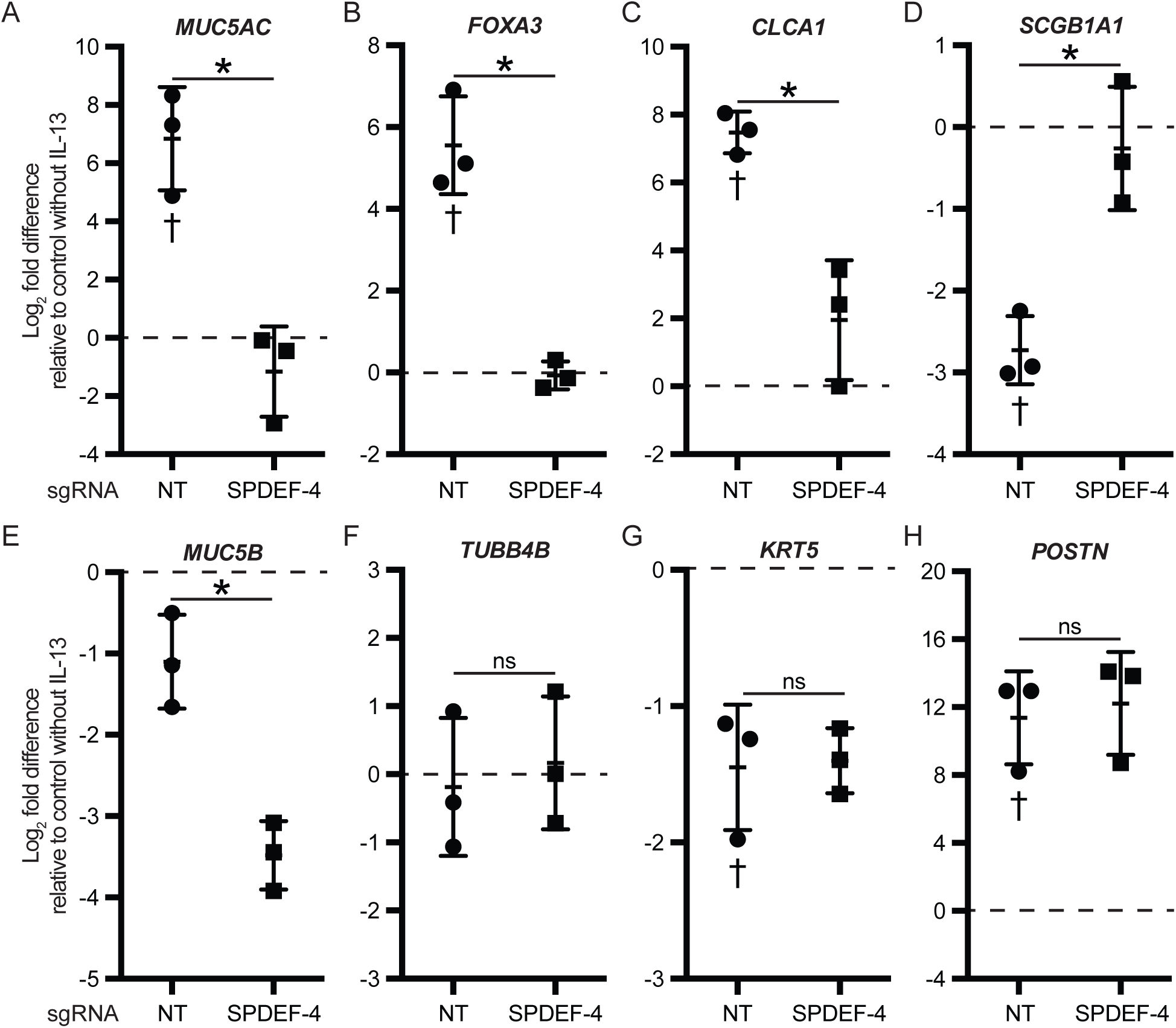
*SPDEF-*dependent and *SPDEF-*independent effects of IL-13 on gene expression in HBECs. Primary HBECs from 3 donors were electroporated twice with RNPs containing non-targeting control sgRNAs (NT) or SPDEF-4 sgRNA and cultured at ALI with IL-13 stimulation for the last 7 d of culture. Gene expression changes were measured using quantitative RT-PCR. Values are log_2_ fold differences relative to cells electroporated with NT control sgRNAs and cultured without IL-13 stimulation (dotted line). *, *P* < 0.05; ns, not significant for comparison between IL-13-stimulated NT and SPDEF-4 sgRNA by Student’s *t* test. †, *P* < 0.05 for comparison between unstimulated controls and IL-13-stimulated HBECs electroporated with NT control sgRNAs by Student’s *t* test.

We also used transcriptional profiling to examine whether *SPDEF* targeting had effects outside of the secretory cell lineage. *SPDEF* targeting had no effects on expression of the ciliated cell marker tubulin beta 4B class IVb (*TUBB4B*) or the basal cell marker keratin 5 (*KRT5*) (Figure 3F and G). We also examined expression of periostin (*POSTN*), which is highly induced by IL-13 and is expressed predominantly in basal cells (24). Induction of *POSTN* was unaffected by targeting *SPDEF* (Figure 3H), which indicates that the targeting protocol did not have global effects on IL-13 signaling outside of the secretory cell lineage.

### Targeting *SPDEF* prevents IL-13-induced goblet cell metaplasia and mucostasis

We used histologic staining and immunofluorescence to examine effects of RNP-based gene targeting on the epithelium (Figure 4). As expected, IL-13 induced a marked change in the appearance of the epithelium, with the appearance of a large number of AB-PAS and MUC5AC-stained goblet cells. Electroporation with rCas9 and a non-targeting control sgRNA had no obvious effects on the appearance of either untreated or IL-13-treated cultures. In contrast, use of a *SPDEF*-targeting sgRNA resulted in dramatic protection from the effects of IL-13 on mucous metaplasia and MUC5AC staining while the population of acetylated alpha-tubulin containing ciliated cells was maintained.

**Figure 4.**
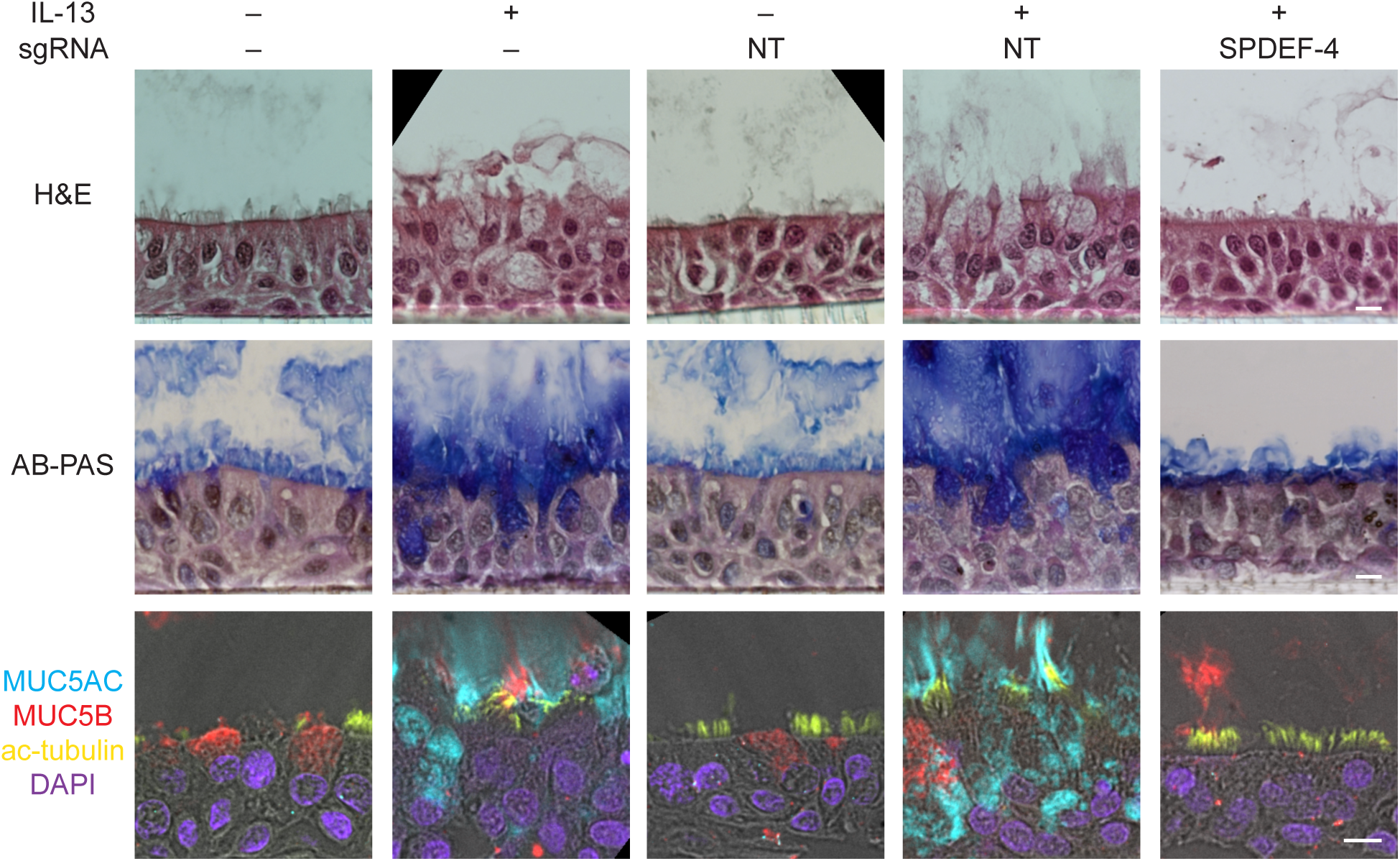
*SPDEF* targeting blocks IL-13–induced mucous cell metaplasia. HBECs were cultured at ALI for 23 d without prior electroporation (sgRNA –) or after electroporation with non-targeting sgRNAs (NT) or SPDEF-4 sgRNA. Some cultures were stimulated with IL-13 for the final 7 d as indicated. (A) Hematoxylin and eosin (H&E) staining. Alcian blue-periodic acid Schiff (AB-PAS), pH, 2.5, staining for mucopolysaccharides. (C) Immunofluorescence staining for MUC5AC (cyan) and MUC5B (red) mucins and the ciliated cell marker acetylated alpha-tubulin (ac-tubulin, yellow). Nuclei were stained with DAPI (purple). Scale bars: 10 *µ*m.

We previously showed that IL-13-induced changes in mucin composition and organization result in epithelial mucus tethering and impaired mucociliary clearance as measured by transport of fluorescent microspheres in HBEC ALI cultures (23). We investigated whether rCas9 and sgRNA-mediated gene targeting could be used to identify molecular mechanisms required for this critical function of HBECs (Figure 5). IL-13 stimulation of cells treated with rCas9 and a non-targeting control sgRNA led to a marked reduction in microsphere transport speed, as previously reported for untransfected cells (23). After targeting *SPDEF*, IL-13-stimulated cultures had transport speeds that were higher than those found in cells electroporated with control sgRNA and treated with IL-13 and were not significantly different from transport speeds found in untreated control cultures. We conclude that IL-13-induced mucostasis is SPDEF-dependent, and that RNP-directed gene targeting enables evaluation of the roles of specific genes in HBEC function.

**Figure 5.**
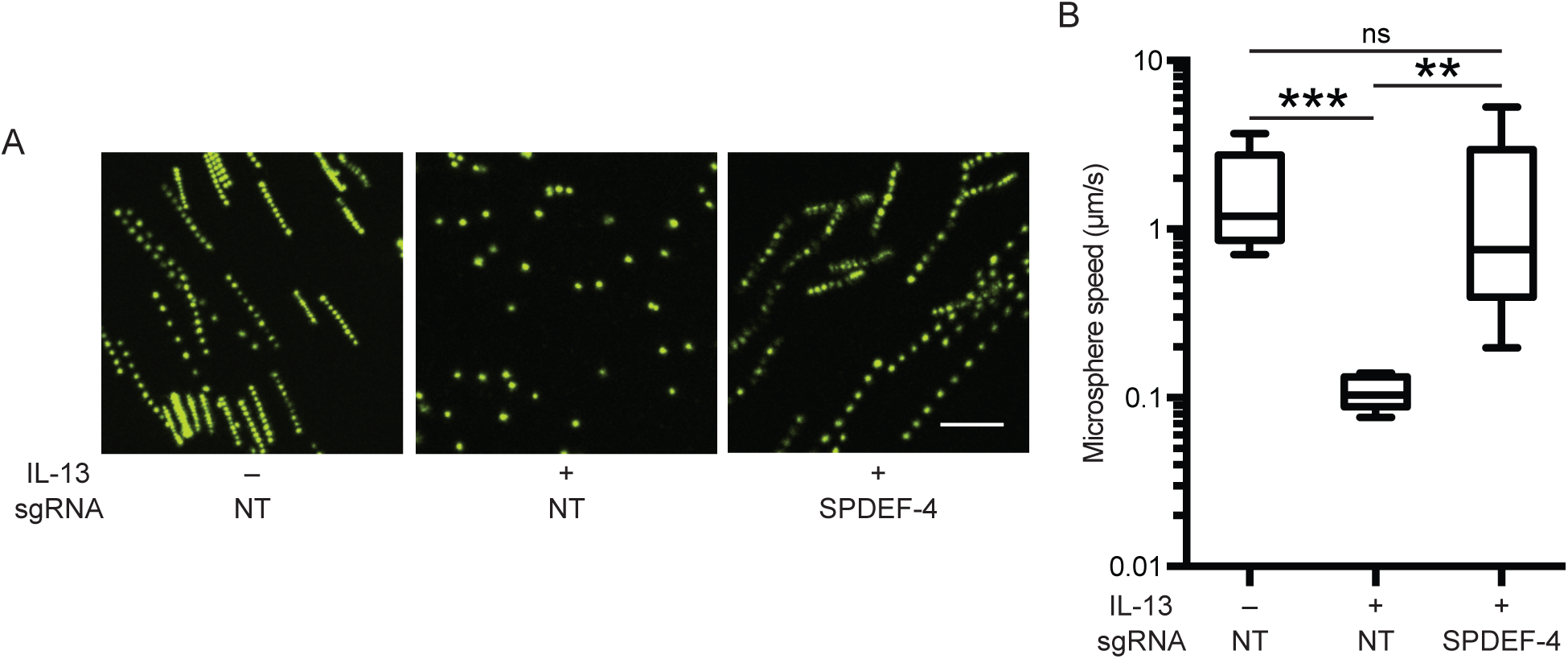
*SPDEF* targeting prevents IL-13–induced mucostasis. Mucociliary transport rates were determined from trajectories of fluorescent microspheres placed on gels atop unstimulated and IL-13–stimulated ALI-cultured HBE cells electroporated with either non-targeting (NT) or *SPDEF*-targeting (SPDEF-4) sgRNAs. (A) Superimposition of 10 images acquired at 1-s intervals. Scale bars: 50 *µ*m. (B) Microsphere speeds determined from three donors, three wells per donor, three fields per well (total of 27 fields per condition). Values represent median microsphere speed for each field. Boxes extend from the 25^th^ to the 75^th^ percentile, the horizontal line within the box indicates the mean, and the whiskers represent minimum and maximum values. **, *P* < 0.01; ***, *P* < 0.001; ns, not significant by the Games-Howell test.

## Discussion

We optimized non-plasmid, non-viral direct delivery of RNPs to HBECs and used this approach to show that the goblet cell transcription factor SPDEF is required for IL-13-induced mucostasis. The methods we developed should be broadly applicable for mechanistic experiments in the widely used HBEC ALI culture system, which has unique strengths for studies of airway epithelial function in human health and in airway diseases. Using these methods, we demonstrated that SPDEF’s central role in goblet cell differentiation is conserved between mouse and human and that disruption of *SPDEF* prevents IL-13-driven changes in secretory cell transcripts. Some effects of IL-13, including its effects on expression of *Muc5b/MUC5B* are not conserved between mouse and human, and the use of the HBEC gene targeting system allowed us to show that *SPDEF* disruption does not restore *MUC5B* expression in IL-13 treated human cells to unstimulated levels. Since the HBEC ALI culture system is well suited for functional studies, we were also able to show that SPDEF is required for IL-13-induced mucostasis, an important contributor to airway obstruction, morbidity, and mortality in asthma.

We identified several factors that affected gene targeting efficiency in HBECs. By varying the amounts of sgRNA and rCas9, the sgRNA sequence, and the number of electroporations, we were able to obtain *SPDEF* targeting efficiencies that approached 100% without the need for antibiotic selection. For the set of four *SPDEF* sgRNAs we tested, targeting efficiency in an immortalized cell line (BEAS-2B) was predictive of efficiency in HBECs. Furthermore, targeting efficiency resulting from direct delivery of RNPs by electroporation also predicted efficiency of targeting by lentiviruses expressing the same gRNA sequence. Although these experiments were limited to a small number of gRNAs targeting the same gene, the results suggest that testing gRNA sequences by direct delivery to cell lines provides an efficient path for selecting optimal gRNA sequences for subsequent use.

Compared with lentiviral delivery, direct delivery of RNPs by electroporation offers several advantages. With direct delivery of RNPs, there is no requirement for plasmid construction, virus production, or antibiotic selection of transduced cells, and there are no concerns about potential effects of viral infection or prolonged expression of Cas9 and gRNAs. By varying delivery conditions, we were able to precisely control targeting efficiency to examine the relationship between the expression of the targeted gene (e.g., *SPDEF*) and the downstream effects of targeting the gene (e.g., *SPDEF*) and the effects of gene targeting (e.g., reduction in *MUC5AC* expression). Although the limited duration of exposure to Cas9 and gRNA might be beneficial in reducing off-target effects, we did not directly assess off-target effects in our experiments. However, the finding that *SPDEF* targeting efficiency was highly correlated with downstream changes in *MUC5AC* expression across the set of four sgRNAs provides confidence that these changes did not result from off-target effects.

The direct delivery method also simplifies targeting of multiple sequences in the genome. We showed that targeting of two sites in the same region led to efficient deletion of larger segments of genomic DNA (>0.5 kb). This approach could be useful for analyzing the functions of regulatory regions, where it may be challenging or impossible to disrupt function with short insertions or deletions produced by individual gRNAs. Although not tested in these experiments, we expect that use of multiple sgRNAs targeting different genes in a single electroporation or in sequential electroporations will allow for efficient analysis of epistatic interactions in HBECs. Despite the advantages of direct delivery of RNPs, lentiviral delivery may be preferred for experiments involving very large numbers of cells, where cost of sgRNA and rCas9 may be an issue, or for testing large libraries of gRNA sequences in pooled screens (25).

We chose to target *SPDEF* based on prior work in transgenic mice that demonstrated a critical role of this transcription factor in goblet cell differentiation (15). In mouse and human systems, IL-13 produced by type 2 lymphocytes and innate lymphoid cells drives an increase in *Spdef/SPDEF* expression and a large change in the secretory cell population, with a decrease in Club cells and an increase in mucus-producing goblet cells (26, 27). IL-13 increases expression of the major mucin gene *Muc5ac/MUC5AC* in both mice and humans (22, 28). In mice, IL-13 also increases expression of the other major airway mucin gene, *Muc5b* (22). However, in experiments with the HBEC human model system, IL-13 tends to decrease *MUC5B* expression (23). This is likely relevant to asthma, since individuals with type 2 high asthma have reduced *MUC5B* expression as well as increased *MUC5AC* expression (29). Experiments in human cell lines found a role for SPDEF in regulation of *MUC5AC* expression (15, 30), but these systems do not provide effective models for studying important aspects of bronchial epithelial cell differentiation or mucociliary function. Targeting *SPDEF* in HBECs allowed us to investigate the role of this transcription factor in a differentiated model of the bronchial epithelium.

We found that targeting *SPDEF* in HBECs led to a proportional reduction in IL-13-induced expression of *MUC5AC*. Using conditions that led to highly efficient targeting showed that *SPDEF* was also required for expression of other genes that are characteristic of goblet cells, including *FOXA3* and *CLCA1*, and for reduced expression of the Club cell gene *SCGB1A1 (CCSP).* Unlike these other changes in secretory cell gene expression, IL-13-induced suppression of *MUC5B* was not dependent upon *SPDEF.* In fact, *MUC5B* levels were significantly lower when IL-13-stimulated cells had been depleted of SPDEF. This indicates that IL-13-stimulated decreases in MUC5B, which are seen in type 2 high asthma but not in mouse models, are mediated in a SPDEF-independent manner.

We found that *SPDEF* is required for IL-13 induced mucostasis. We previously showed that IL-13-induced changes in mucus production and organization result in tethering of MUC5AC-rich mucus domains to the airway epithelium, which dramatically impairs mucociliary clearance (23). The ability to efficiently target genes in the HBEC ALI culture system without apparent non-specific effects on cell differentiation or mucociliary function allowed us to test the requirement for *SPDEF* in IL-13 induced mucostasis. Our results suggest that SPDEF could be a valuable target for therapies designed to decrease mucus production and improve mucociliary clearance in asthma or other airway diseases. However, given the important role of MUC5B in airway homeostasis and protection from infection (31), the finding that *SPDEF* targeting exacerbates rather than reverses IL-13-induced decreases in *MUC5B* expression suggests a potential limitation of this approach.

## Acknowledgements

The authors thank Michael Matthay for supplying human bronchial specimens and Jane Gordon (UCSF Laboratory for Cell Analysis), Kari Herrington (UCSF Nikon Imaging Center), Theodore Roth, Alexander Marson, and Olivier Le Tonqueze for technical advice and assistance.

