## Supplementary material for "Efficient RNP-directed human gene targeting reveals SPDEF is required for IL-13-induced mucostasis": Online Data Supplement

### Supplementary Materials and Methods

#### Cells

The UCSF Committee on Human Research approved the use of HBECs isolated from lungs not used for transplantation. Written consent was not required since materials were leftover clinical samples obtained from de-identified individuals. Prior to electroporation, HBECs were seeded onto 10-cm dishes coated with human placental collagen (HPC; Sigma-Aldrich, St. Louis, MO) and propagated in BEGM (Lonza, Walkersville, MD) with 10 μM Y-27632 (Enzo Life Sciences, Farmingdale, NY) (1). For experiments with BEAS-2B cells (ATCC, Manassas, VA), cells were cultured in 1:1 DMEM/F12 medium with 10% FBS (Thermo Fisher Scientific, Waltham, MA).

**Lentiviral delivery of sgRNA and Cas9**

gRNA sequences (Table E1) were used to design a pair of complementary 5′-phosphorylated oligonucleotides (5′-$PO_4$-CACCNNNNNNNNNNNNNNNNNNNN-3′ and 5′-$PO_4$-AAACNNNNNNNNNNNNNNNNNNNN-3′; where Ns indicate sequences that vary according to the gRNA). gRNA oligonucleotides (IDT, San Jose, CA) were annealed and cloned into the LentiCRISPRv2 plasmid (2) (Addgene, Watertown, MA) digested with BsmBI (New England Biolabs, Ipswich, MA). Lentivirus was produced by transfecting LentiCRISPRv2-gRNA plasmids and three lentivirus-packaging plasmids (pMDL, pVSV-G, and pRSV-Rev) into HEK293T cells using TransIT-293 Transfection Reagent (Mirus Bio, Madison, WI). Harvested lentivirus was concentrated 100-fold using Lenti-X Concentrator (Takara Bio, Mountain View, CA). After propagation in HPC-coated 10-cm dishes and passaging to 6-well dishes, HBECs were transduced with lentivirus, and transduced cells were maintained in BEGM with 10 μM Y-27632 and 1 μg/mL puromycin (Thermo Fisher Scientific) for 3 d for selection before passaging to Transwell inserts (Corning, Corning, NY) for ALI culture.

**Quantification of MUC5AC-expressing cells**

Accumulated mucus was removed from the apical surface of HBEC cultures with 10 mM dithiothreitol (DTT; Enzo Life Sciences, Farmingdale, NY) in PBS for 10 min at 37 °C. HBECs were subsequently washed with PBS three times, trypsinized with 0.25% trypsin-EDTA (Thermo Fisher Scientific) for 10-15 min at 37 °C, neutralized, fixed in 4% paraformaldehyde (Thermo Fisher Scientific) for 10 min at 4 °C, and washed with PBS. Fixed HBECs were then incubated for 30 min at 4 °C in PBS containing 5% normal goat serum (Jackson ImmunoResearch Laboratories, West Grove, PA) to block non-specific staining and 0.2% saponin (Sigma-Aldrich) for permeabilization, prior to a 1 h incubation with mouse monoclonal anti-MUC5AC (45M1;

Novus Biologicals, Centennial, CO) conjugated with Dylight 488. Cells were then washed twice and resuspended in eBioScience Flow Cytometry Staining Buffer (Thermo Fisher Scientific) prior to flow cytometry (FACS Canto II, BD Biosciences, San Jose, CA; data analysis with FlowJo, FlowJo LLC, Ashland, OR). The threshold for MUC5AC-positive cells was established based on staining of IL-13-treated cells with an isotype control antibody (<0.5% of isotype control-stained cells exceeded the threshold).

**Quantification of mRNA transcripts**

mRNAs were measured by quantitative real-time RT-PCR (qRT-PCR). After removing accumulated mucus, RNA was extracted from HBECs using Buffer SKP from the RNA/DNA/Protein Purification Plus Kit (Norgen Biotek) according to the manufacturers' protocol. RNA was reverse-transcribed using SuperScript III First-Strand Synthesis System (Thermo Fisher Scientific), and the resulting cDNA was analyzed by qRT-PCR (PowerUp SYBR Green, Thermo Fisher Scientific; primer sequences in Table E2). The mean value of three technical replicates was used for analysis. mRNA levels were normalized to *GAPDH* levels and comparisons were made using the using $\Delta\Delta C_t$ method (3).

**Histologic staining and immunofluorescence**

Paraffin-embedded microscopic sections of HBEC culture Transwell inserts were prepared as described previously (4). Hematoxylin and eosin (H&E) staining was done as described previously (5). Alcian blue-periodic acid Schiff (AB-PAS) staining was done using Alcian Blue pH 2.5 Periodic Acid Schiff Stain (Thermo Fisher Scientific) and following manufacturer's instructions. Immunofluorescence staining was done with following primary antibodies: mouse monoclonal anti-MUC5AC (45M1; Thermo Fisher Scientific; 1:150), rabbit polyclonal anti-MUC5B (H-300, sc-20119; Santa Cruz Biotechnology, Dallas, TX; 1:150), and mouse monoclonal anti acetylated alpha tubulin (6-11B-1, sc-23950; Santa Cruz Biotechnology; 1:200). After washing, slides were incubated with appropriate secondary antibodies (Rhodamine goat anti-mouse, Alex Fluor 647 goat anti-rabbit, and Alexa Fluor 488 goat anti-mouse; Jackson ImmunoResearch Laboratories) at 1:200 dilution for 1 h. 4′,6-diamidinio-2-phenylindole (DAPI) was used to stain nuclei. Immunofluorescence mages were acquired using a Yokagawa CSU22 spinning-disk confocal microscope connected to a Nikon Ti-E (Nikon Imaging Center, UCSF). Slides were placed on the microscope stage and fluorescence and brightfield images were acquired using a 20× objective. Identical acquisition settings were used throughout each experiment.

**Measurement of mucociliary transport**

Methods for measuring mucociliary transport were adapted from previous reports (4, 6–8). 2-μM yellow-green (505/515) fluorescent microspheres (Thermo Fisher Scientific) were applied to the apical surface of HBEC cultures with intact mucus gels and allowed to disperse for 10 min. Cultures were then transferred to an optical cell dish (MatTek, Ashland, MA) and placed on the stage of a spinning disc confocal microscope under a dry 10× objective at 37 °C with perfluorocarbon (Sigma-Aldrich). Transport of fluorescent microspheres was imaged in the plane of the gel by recording sequential images every 1 s over 1 min. Images were analyzed using the TrackMate plugin in Fiji. Median microsphere speed was determined for each of three fields from each of three wells for each of three donors.

**Statistics**

Student's *t* test was used to compare qRT-PCR $\Delta\Delta C_t$ values. The Games-Howell test was used to compare particle compare tracking speeds. A *P* value of less than 0.05 was considered statistically significant.

**Table E1.** gRNA sequences

| Name | gRNA target DNA sequence (5′-3′) |
|---|---|
| NT-1 | GACGACTAGTTAGGCGTGTA |
| NT-2 | GGCCAAACGTGCCCTGACGG |
| SPDEF-1 | TAGGTGGCTCAACAAGGAGA |
| SPDEF-2 | GAGCTGGACCGACAGCGAGG |
| SPDEF-3 | GGAGAGCTGGACCGACAGCG |
| SPDEF-4 | ATGAAGCGGCCATAGCTGTG |
| KLF5 | GTGTGTTACGCACGGTCTCT |
| c1 | AAAGTCTGAGGGTATGTACA |
| c2 | CAGACCCTCTTGAGTCACTT |
| s1 | GTGATGTCGGGGAAATGGTT |
| s2 | GTTCTCTTGGCAATCAGACA |
| s3 | AGTCTTGCATGCAGGCCCTT |
| s4 | GAGTCTTGCATGCAGGCCCT |

**Table E2.** PCR, sequencing, and RT-PCR primers

| Name | Primer sequence (5′-3′) |
|---|---|
| SPDEF-PCR.f | CTCACTTGGCAAGAGCATCC |
| SPDEF-PCR.r | CCATGTCAGATGTCCTCATCTG |
| SPDEF-seq | TGTCCCATGAGAGCTGCATA |
| KLF5-PCR.f | CAGGGAATCCTGCTTGTACAG |
| KLF5-PCR.r | TAAGTGGCCTGTTGTGGAAG |
| KLF5-seq | ATCCGGTGTATTCAGTAGCT |
| D-PCR.f | TCACAGTGTCTCAGAACTAAGGG |
| D-PCR.r | TCCTGTAAACTTGTGTGGCC |
| q.MUC5AC.f | CAACATCAGGAACAGCTTCGA |
| q.MUC5AC.r | GAGCACCAGTGCTGAGCAT |
| q.FOXA3.f | TGGCCGAGTGGAGCTACTAC |
| q.FOXA3.r | GGATTCAGGGTCATGTAGGAGT |
| q.CLCA1.f | TCCAATGGAAGAATACAAGCAG |
| q.CLCA1.r | AATGTGCATCTTTTGGTGTAACAG |
| q.SCGB1A1.f | GCTGAAGAAGCTGGTGGACAC |
| q.SCGB1A1.r | TGCTAATTACACAGTGAGCTTTGG |
| q.MUC5B.f | GCTCCAAGGCCATCAAGCT |
| q.MUC5B.r | GGTCTCGATGACCAGGAAGATC |
| q.KRT5.f | AGCAGTGGTACGCTTGTTGATT |
| q.KRT5.r | GCCTGGACTCAGAGCTGAGAA |
| q.TUBB4B.f | ATCTTCCGGCCGGACAAC |
| q.TUBB4B.r | CATCCAGCACCGAGTCCA |
| q.POSTN.f | GACCGTGTGCTTACACAAATTG |
| q.POSTN.r | AAGTGACCGTCTCTTCCAAGG |
| q.GAPDH.f | CACATCGCTCAGACACCAT |
| q.GAPDH.r | GGCAACAATATCCACTTTACCAG |
